## Supplementary figures and images for "Intestinal permeability and peripheral immune cell composition are altered by pregnancy and adiposity at mid- and late-gestation in the mouse"

### https://docs.google.com/document/d/1dzLxNeMcxpmDaGCMpAR9Oc6AWY-6uiRG9VtnVk7HogU/edit

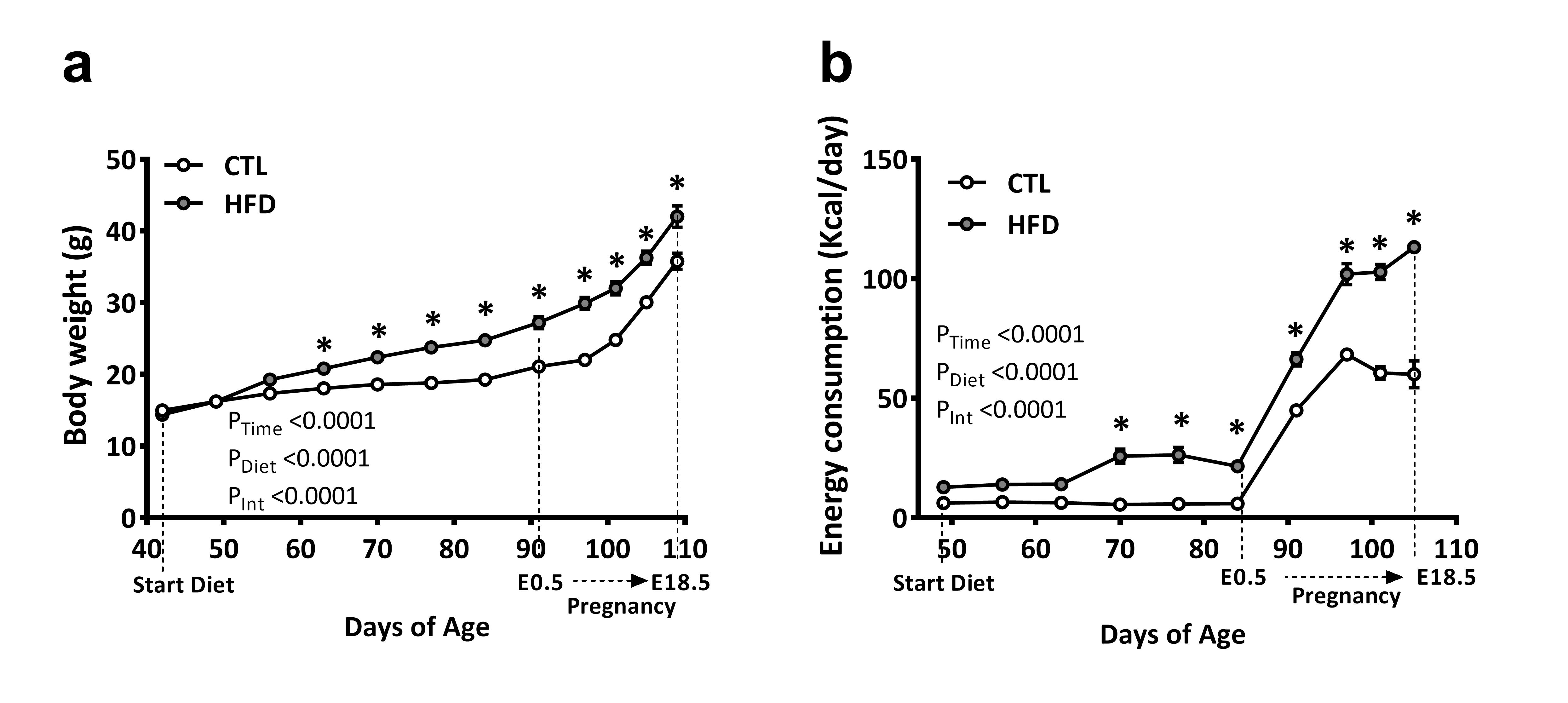

### https://docs.google.com/document/d/1dzLxNeMcxpmDaGCMpAR9Oc6AWY-6uiRG9VtnVk7HogU/edit

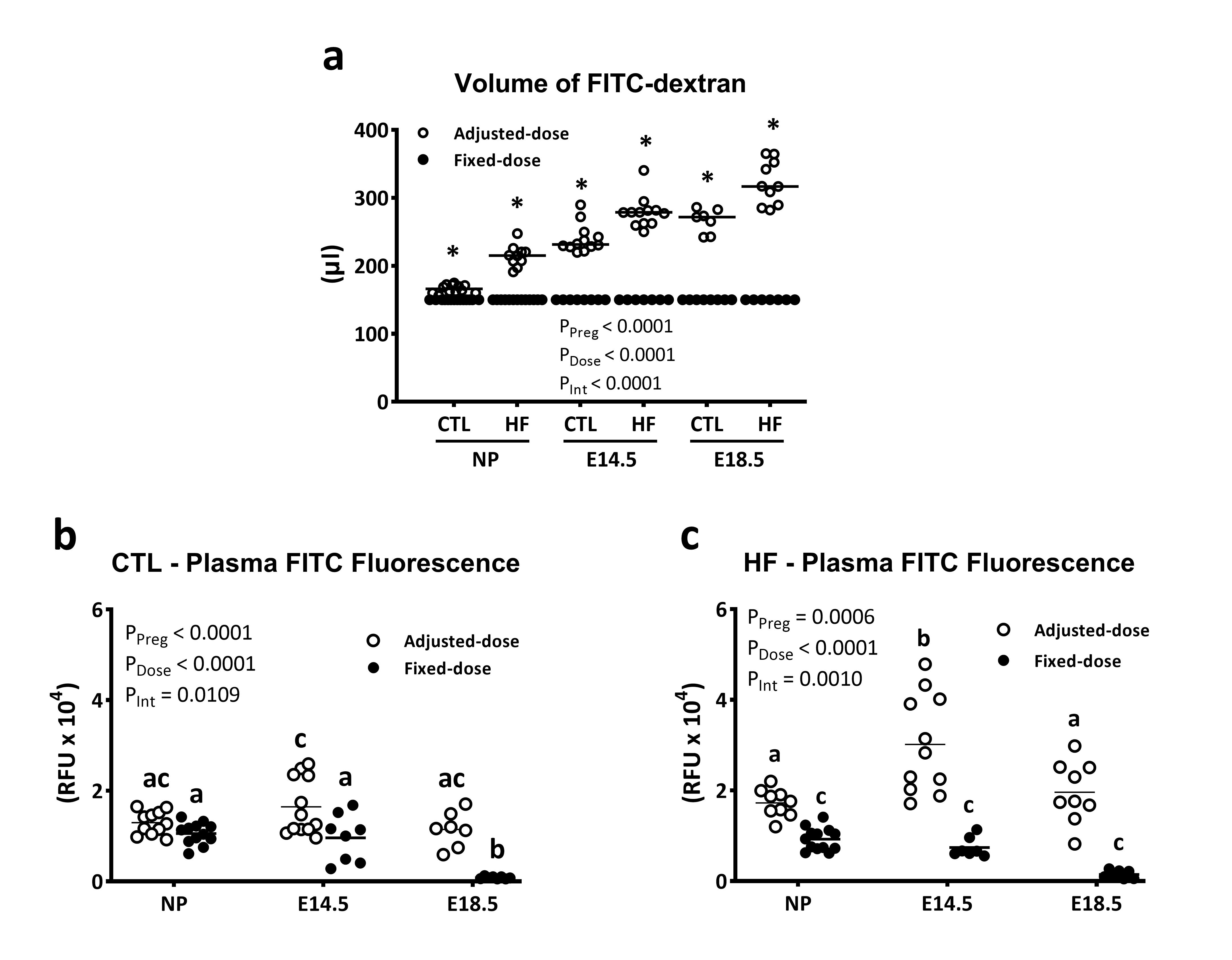

### https://docs.google.com/document/d/1dzLxNeMcxpmDaGCMpAR9Oc6AWY-6uiRG9VtnVk7HogU/edit

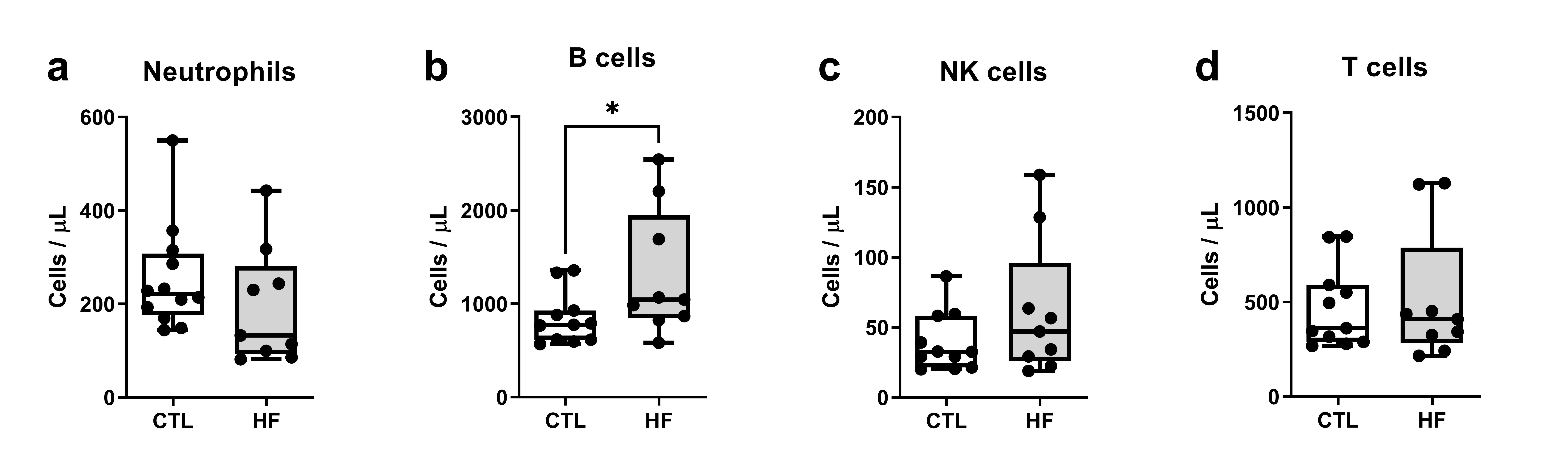
